# Informative Neural Codes to Separate Object Categories

**DOI:** 10.1101/2020.12.04.409789

**Authors:** Mozhgan Shahmohammadi, Ehsan Vahab, Hamid Karimi-Rouzbahani

**Affiliations:** Islamic Azad University, Central Tehran Branch, Tehran, Iran; Islamic Azad University, Qazvin Branch, Qazvin, Iran; Macquarie University, Sydney, Australia

**Keywords:** Object Category Recognition, Feature Extraction, Brain Decoding, Classification

## Abstract

In order to develop object recognition algorithms, which can approach human-level recognition performance, researchers have been studying how the human brain performs recognition in the past five decades. This has already inspired AI-based object recognition algorithms, such as convolutional neural networks, which are among the most successful object recognition platforms today and can approach human performance in specific tasks. However, it is not yet clearly known how recorded brain activations convey information about object category processing. One main obstacle has been the lack of large feature sets, to evaluate the information contents of multiple aspects of neural activations. Here, we compared the information contents of a large set of 25 features, extracted from time series of electroencephalography (EEG) recorded from human participants doing an object recognition task. We could characterize the most informative aspects of brain activations about object categories. Among the evaluated features, event-related potential (ERP) components of N1 and P2a were among the most informative features with the highest information in the Theta frequency bands. Upon limiting the analysis time window, we observed more information for features detecting temporally informative patterns in the signals. The results of this study can constrain previous theories about how the brain codes object category information.

## 1 Introduction

Object recognition algorithms are widely used in computer vision applications such as image annotation (Vedaldi & Lenc, 2015), face recognition (O’Toole et al., 2018), face detection (Kalinovskii, n.d.) and object tracking (El-Shafie et al., 2019). They are categorized into one of the two categories of “machine learning” approaches and “deep learning” approaches. In the machine learning approaches, one needs to first define a set of statistical/topographical/mathematical features (e.g. Haar features, Scale-invariant feature transform (Lowe, 1999), histogram of oriented gradients (Dalal & Triggs, 2005)), extract them from object images and then apply a machine-learning classifier such as support-vector machine (SVM) to categorize the objects into their semantic categories (e.g. car, animal, human, etc.). The deep learning approaches (including Region Proposals (Girshick et al., 2014) and Single Shot Multibox Detectors (Liu, 2016)) which generally use convolutional neural networks, on the other hand, do not need the features to be defined prior to classification; they do the feature extraction and classification within the same architecture. Interestingly, the deep learning approaches, which are in many ways inspired by how the human visual system processes object features and performs recognition (Karimi-Rouzbahani et al., 2017c; Khaligh-Razavi & Kriegeskorte, 2014; Kheradpisheh et al., 2016), have outperformed most of the machine-learning approaches in the past eight years (Krizhevsky et al., 2012). In fact, the object recognition performance surged upon the introduction of second generation of convolutional neural networks, namely AlexNet, in 2012 (Krizhevsky et al., 2012), after more than a decade of slow improvements. Since then, one of the main approaches that researchers have been pursuing to improve the recognition performance, has been to understand how the human brain performs object recognition (Federer et al., 2020). These insights will in turn allow us to develop more brain-like object recognition algorithms which can potentially reach human-level performance.

To understand how the human brain performs object recognition, the first step is to understand how we can ‘read out’ or ‘decode’ the codes that the brain generates for each object category. Therefore, we need to quantitatively compare large sets of features of the brain activity to characterize the informative ones for decoding. This is important, as it will constrain the understudied parameters of the available brain-inspired deep learning algorithms for improved recognition performance. Here we evaluate the information content of large-scale feature set of electrical brain activity (EEG), recorded while humans performed a simple object recognition task. To that end, we use multivariate pattern decoding (or simply decoding) to capture the most subtle differences in the patterns of neural activity across different object categories (Hebart & Baker, 2018).

During the past decade, a number of DECODING studies have evaluated the information content of brain activity as recorded through EEG to decode the category of the presented objects from brain activity (Kaneshiro et al., 2015; Karimi-Rouzbahani et al., 2017a; Simanova et al., 2010). For example, earlier studies have shown that the mean amplitudes of brain activations such as P1, N1, P2a and P2b, which indicate average amplitudes in specific time windows of signal time series, between 100 to 300 ms object presentation time, could be used to reliably discriminate object categories (Chan et al., 2011). Later investigations implemented a classifier of Linear Discriminant Analysis (LDA) and showed the components were so informative that allowed the discrimination of up to four object categories (Wang et al., 2012). Other studies investigated more complex ways in which information coding could happen in the brain such as frequency coding (Wang et al., 2018), signal power coding (Majima et al., 2014; Miyakawa et al., 2018; Rupp et al., 2017), coding based on within-trial correlation (Karimi-Rouzbahani et al., 2017a; Majima et al., 2014), and non-linear statistical features such as signal Kurtosis, Hjorth complexities, fractal dimensions, etc. which generally represent the randomness of the signals can be used to discriminate visual object categories (Stam, 2005; Sweeti et al., 2018; Torabi et al., 2017).

While all of the above-mentioned studies have been able to categorize visual object categories using their proposed signal features, their results lack the generalizability to other studies. We still do not know which of these features carry more object category information. This is important since, while the secondary attributes of the neural code may allow for the classification of categories, they still fall behind the most informative features, which might be available but overlooked. Therefore, we need to compare as many features as possible on a well-controlled dataset to assure which feature is the most informative. In fact, while several of the above-mentioned studies compared multiple features, most of the proposed features were used in separate studies, sometimes causing discrepancy in the results. For instance, one study showed that object categories could be discriminated from signal phase patterns in low frequency bands but not from signal power (Behroozi et al., 2016), a finding that was later repeated by another group, however, this time in a higher frequency band (4-8 Hz vs 1-4 Hz; (Wang et al., 2018)). To overcome this issue, we extracted a large set of 25 features, proposed in previous studies, from a generalizable object recognition dataset and evaluated their performance in providing object category information.

## 2 Methods

### 2.1 Dataset

We used a previously published EEG dataset for this study. This dataset has been used in a time-resolved object categorization decoding study and has provided significant information about object categories using multivariate pattern analysis (Karimi-Rouzbahani et al., 2019).

#### *Image set and* task

EEG signals were collected from 10 human participants, while they performed an object recognition task as shown in Fig. 1B. Basically, they were presented with (48 unique) images from one of four categories of objects including animals, fruits, objects and human faces as shown in Fig. 1A. Participants were instructed to press the spacebar of a computer if the presented object was from the category indicated (for 5000 ms) at the beginning of each blocks of trials and not if it was from other categories. Each block of trials included 12 image presentations and each image was presented for 900 ms followed by 800 ms of inter-stimulus-interval (Fig. 1B). Each unique stimulus was presented 6 times to increase the signal-to-noise ratio for analyses.

**Fig. 1.**
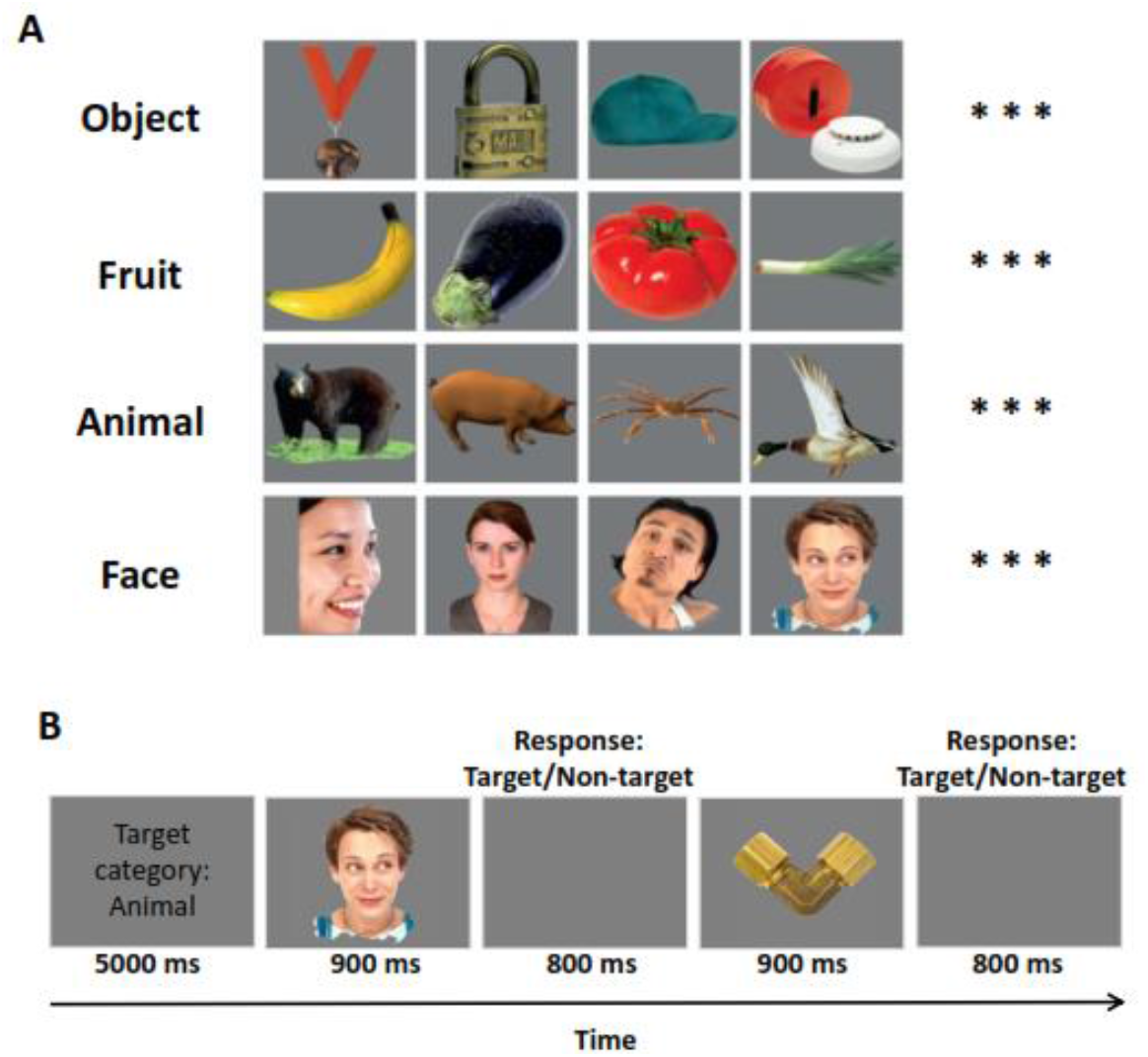
Image set (obtained from Kiani et al., 2007 with permission; A) and the object recognition paradigm (B). A, the image set included 4 categories of images with Object, Fruit, Animal and Face categories each including 12 exemplars (4 shown here). Images were 256*256 color images with balanced average luminance and contrast across categories. B, Subjects were supposed to press the spacebar if whenever they saw an image from the cured target category and refrain from response otherwise. They had 800 ms to respond.

#### Signal pre-processing

The brain signals were sampled at 1000 Hz (using a 31-channel amplifier) and only the EEG data for correctly recognized object images (M=94.65%, SD=9.38% across participants) were used for decoding. We band-pass filtered the EEG signals in the range from 0.03 to 200 Hz to remove baseline drift and high-level noise. We also notch-filtered the signals at 50 Hz to remove line noise and removed eye-movement artifacts using Independent Component Analysis (ICA). We split the signals around the time of stimulus presentation from 1 to 1000 ms after the stimulus onset for analysis. All the analyses were done in Matlab with the preprocessing done using the EEGLAB toolbox (Delorme & Makeig, 2004). For detailed explanation of the dataset see (Karimi-Rouzbahani et al., 2019).

### Extracted Features

We extracted a set of 25 features from the signal time series (from 1 to 1000 ms after stimulus onset) ranging from the simple statistical feature of mean to more complex features of fractal dimensions. Here we very briefly explain each feature. For more information, the readers are referred to the relevant references.

#### Mean, Variance, Skewness and Variance

These features are the standard 1^st^ to 4^th^ moments of EEG time series. To calculate these features, we simply calculated the mean, variance, skewness and variance of EEG signals as defined in statistics. We also present the results for the *Baseline*, which refers to the signal mean in the pre-stimulus time span (−200 to 0 ms) and is expected to provide no information (it is a control feature).

#### Median

We also calculated signal’s median as it is less affected by spurious values compared to the signal mean.

#### LZ complexity

We calculated the Lempel-Ziv complexity as an index of signal complexity. This measure, counts the number of unique sub-sequences within the signal time samples, which is first turned into a binary sequence. We used the signal median to make the signal samples binary. Accordingly, the LZ complexity of a time series grows with the length of the signal and its irregularity. See (Ziv & Lempel, 1976) for more details.

#### Higuchi and Katz fractal dimensions

Fractal is an indexing technique which provides statistical information determining the complexity of how data are organized in a time series. Accordingly, higher fractal value, suggests a more complexity and vice versa. In this study, we calculated the complexity of the signals using two methods of Higuchi and Katz, as had been used before for categorizing object categories (Namazi, 2018).

#### Hurst exponent

This measure, quantifies the long-term memory in a time series. Basically, it calculates the degree of dependence among consecutive samples of time series and functions similarly to the autocorrelation function (Racine, 2011).

#### Sample and Approximate Entropy

Entropy measure the level of perturbation in time series. As precise calculation of entropy needs large sample sizes and is also noise-sensitive, we calculated it using two of the most common approaches. Sample entropy is not as sensitive to the sample size and simpler to implement compared to Approximate entropy. Sample entropy, however, does not take into account self-similar patterns in the time series (Richman & Moorman, 2000).

#### P1, N1, P2a and P2b (event-related potential; ERP) components

These signal components refer to signal amplitude (mean value) within well-known prespecified time windows of trial. These are defined as P1 (80 to 120 ms), N1 (120 to 200 ms), P2a (150 to 220 ms) and P2b (200 to 275 ms).

#### Within-trial correlation

This index quantifies the self-similarity of a time series at specific time lags. Accordingly, if a time series has a repeating pattern at the rate of *F* hertz, an autocorrelation measure with a lag of 1/*F* will return a value of 1. However, it would return −1 at the lag of 1/2*F*. It would provide values between −1 and 1 for other cases. More complex signals, would provide values close to 0.

#### Hjorth complexity and mobility

These parameters measure the variation in the signals’ characteristics. Specifically, the complexity measure, calculates the variation in a signal’s dominant frequency, and the mobility measures the width of the signal’s power spectrum (how widely the frequencies are scattered in the signal; (Torabi et al., 2017)).

#### Mean, Median and Average Frequency

These measures, calculate the central frequency of the signal in different ways. Mean frequency is the average of all frequency components available in a signal. Median frequency is the median normalized frequency of the power spectrum of the signal and the average frequency is the number of time the signal time series crosses zero.

#### Spectral Edge Frequency

It indicates the high frequency, below which, *x* percent of the signal’s power spectrum exists. *X* was set to 95% in this study.

#### Signal power, Power and Phase at Median Frequency

Signal’s power (i.e. power spectrum density) was used as a feature. Signal’s power and phase at median frequency have been shown to be informative about object categories (Jadidi et al., 2017; Rupp et al., 2017). Therefore, we also extracted them from signals to evaluate their information content in category decoding.

### Multivariate pattern analysis (decoding)

We used multivariate decoding to measure the level of information contained in each feature about object categories. This method is basically a classification, which is quite common in machine learning, which takes into account the activity from all electrodes in EEG as input features to the classifier. We used an LDA classifier to classify every pair of categories (6 pairs for 4 categories). We used a 10-fold cross-validation procedure in which we trained the classifier on 90% of data and tested them on the left-out 10%. We repeated this procedure 10 times so that each data is used once as a testing data. We did it for every time window, the main frequency bands in EEG (Delta= 0.5-4Hz; Theta= 4-8Hz; Alpha=8-12Hz; Beta=12-16Hz and Gamma=16Hz>).

### Statistical analysis

In order to determine the significance of decoding accuracies with chance-level decoding, we generated 1000 random accuracies using random permutations of class labels. To that end, we randomized the labels of categories and did the classification as if they were true categories. Finally we compared the true distribution with the null distribution and considered the true accuracies as significant if they surpassed 95% of the accuracies from the null distribution. All the p-values were corrected for multiple comparisons across features using Bonferroni correction.

## 3 Results

The decoding results showed that only a few of the features could provide category information with above-chance accuracy (p<0.05; random permutation testing; Fig. 2A). Specifically, Mean, Variance, Higuchi-frct, Hurst-Exp, Hjorth-Mob, Mean-Freq, Med-Freq, within-trial correlation as well as all ERP components showed information for object categories. Importantly, ERP components provided the highest decoding accuracy as reflected in their mean. We also looked at the results from each individual category separately to see if the coding of information was advantageous for any categories (Fig. 2B). We saw the dominance of the same features, with the most pronounced decoding values for the ‘Face’ category. This is consistent with previous finding showing stronger coding of face category information in the human brain compared to other categories of objects (Kaneshiro et al., 2015; Karimi-Rouzbahani et al., 2017a; Petras et al., 2019).

**Fig. 2.**
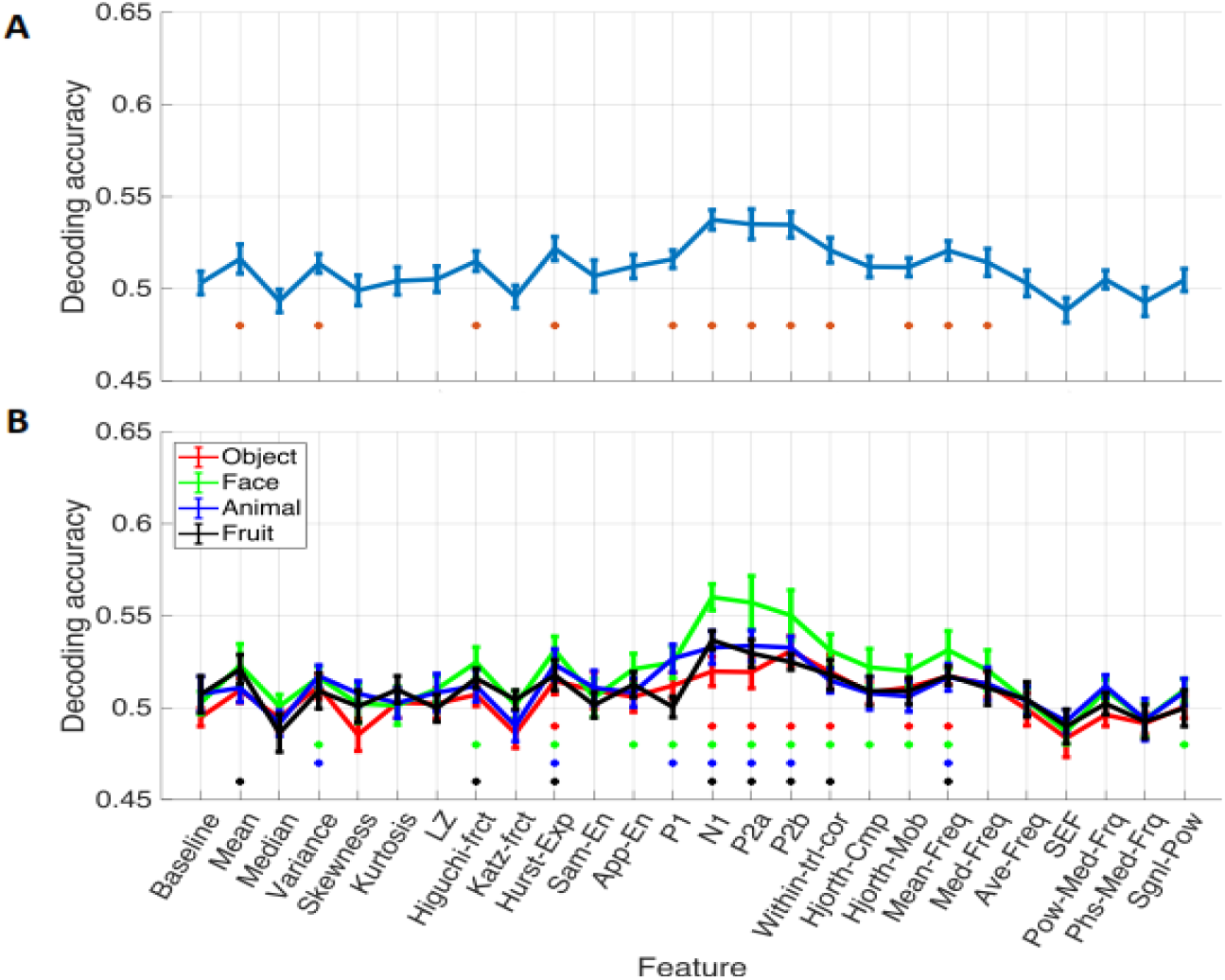
Decoding results for all categories (A) and each individual category (B). A, decoding accuracy reflects the average classification accuracy along with the standard error (SE) across participants (error bars). Colored dots indicate features with significantly (p<0.05; random permutation testing) above-chance accuracy. B, decoding accuracy for each category was calculated as the average of decoding accuracy when classifying the given category from all other categories.

In order to quantify the advantage for the face category, we performed a signed-rank statistical testing to quantitatively compare the decoding results across categories. We observed that, the face category dominated other categories, especially for the ERP components of P1, N1, P2a and P2b (Fig. 3). This is consistent with previous studies which observed the temporally constrained representation of face processing at around 170 ms post-stimulus onset (Nemrodov et al., 2018; Rousselet & Pernet, 2011).

**Fig. 3.**
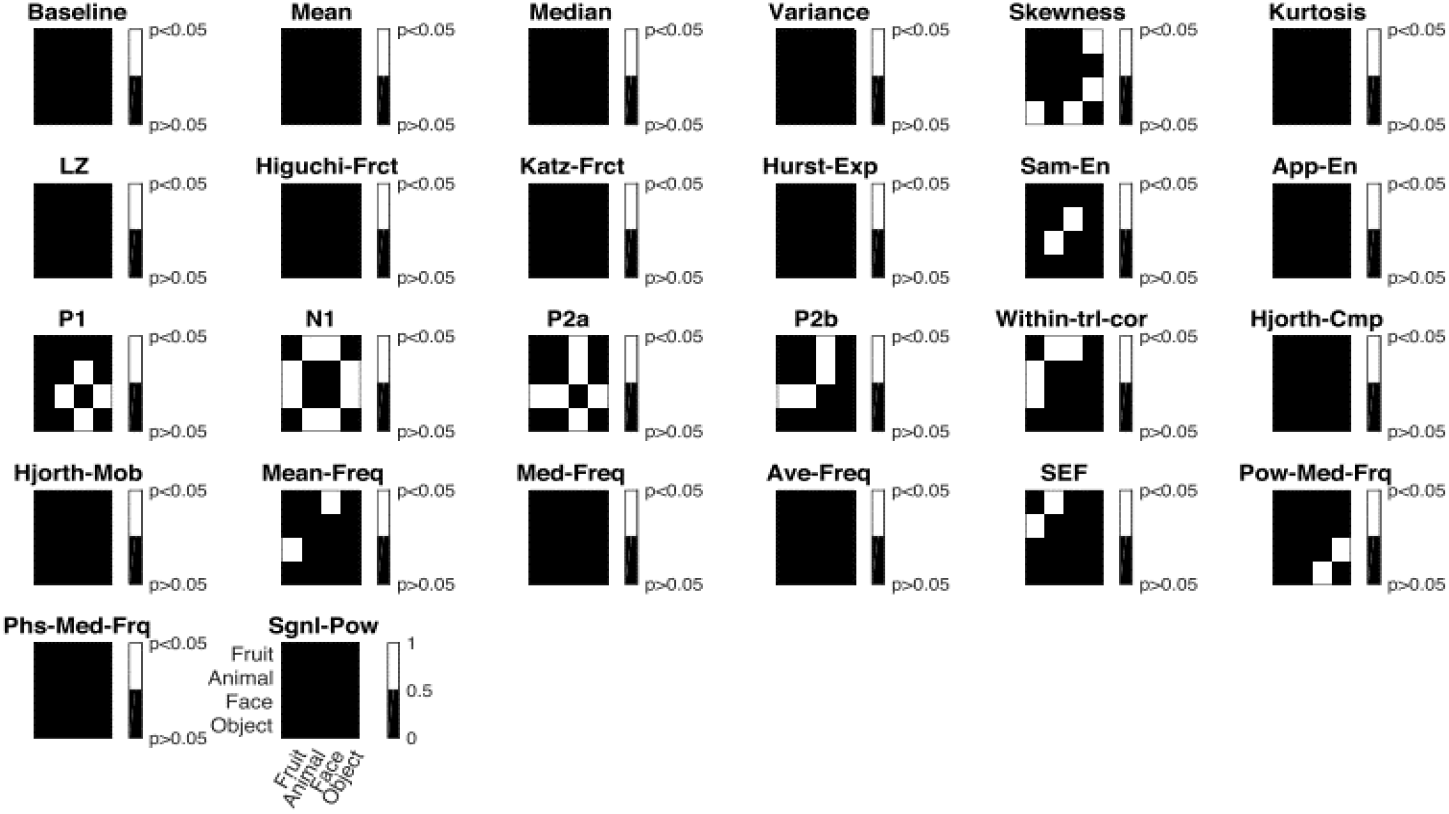
Statistical testing results evaluating the difference between the decoding accuracies obtained for individual object categories (Fig. 2B). Wilcoxon’s signed-rank test was performed between all possible pairs of categories and the p-values were corrected for multiple comparisons across features and combination of categories. Robust results were found supporting the dominance of face category decoding especially for the ERP components.

Motivated by previous studies finding category information in specific frequency bands, we also evaluated the information contents of the well-known EEG frequency bands of Delta= 0.5-4Hz, Theta= 4-8Hz, Alpha=8-12Hz, Beta=12-16Hz and Gamma=16Hz> (Fig. 4). While the Delta band provided above-chance decoding accuracies and seemed to surpass other frequency bands in a few features such as the moment features (e.g. Mean, Variance) as well complexity-measuring features (e.g. Katz-frct), the most information seem to have been represented in the EEG signals in the Theta and Alpha bands. This was more pronounced or the ERP components. The observation that the Theta band dominated other frequency bands can be explained by the processing of visual information in the brain by the feed-forward neural mechanisms (Bastos et al., 2015).

**Fig. 4.**
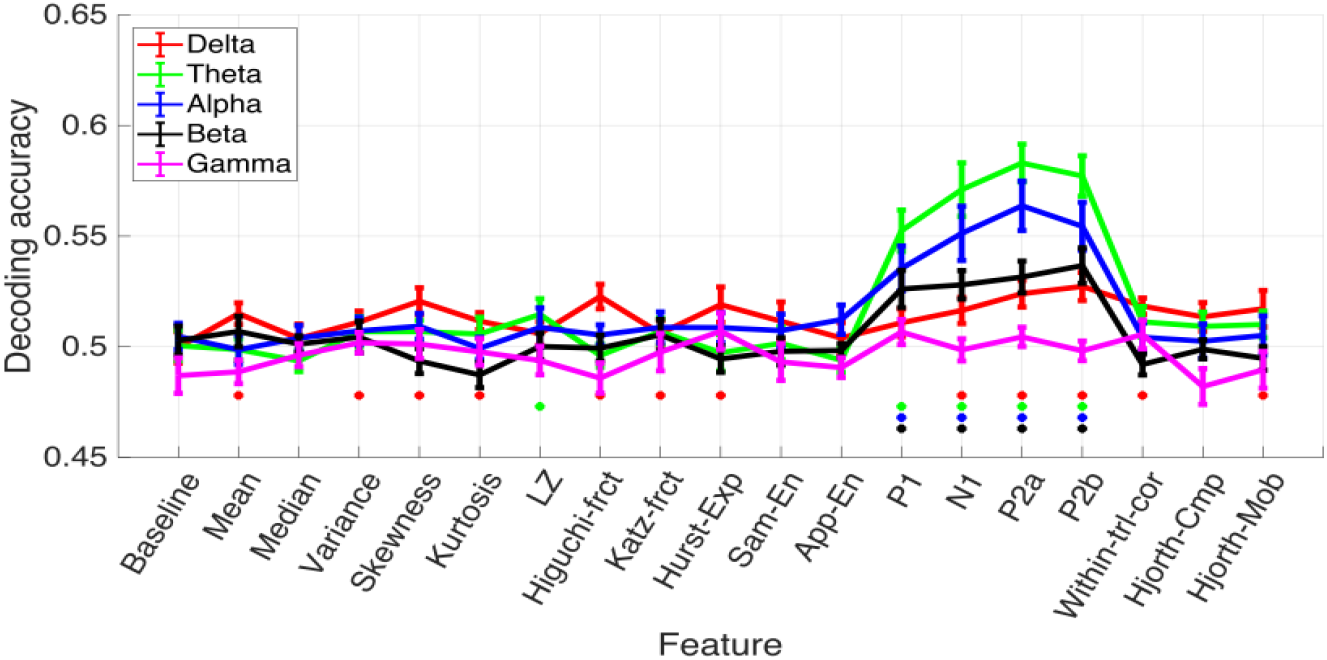
Decoding accuracy obtained for all features (except for frequency-domain features) in the well-established informative EEG frequency bands of Delta, Theta, Alpha, Beta and Gamma. Decoding results reflect the average classification accuracy along with the standard error (SE) across participants (error bars). Colored dots indicate features with significantly (p<0.05; random permutation testing) above-chance accuracy.

The results so far showed that ERP features which were extracted from the signal in the time span from 80 to 275 ms relative to the stimulus presentation onset, provided the most information about visual object categories. This was different to the other features, which were all extracted from the whole trial time span (0 to 1000 ms). These results and previous time-resolved studies (Kaneshiro et al., 2015; Karimi-Rouzbahani, 2018; Karimi-Rouzbahani et al., 2017b; Karimi-Rouzbahani et al., 2020; Karimi-Rouzbahani et al., 2020a) suggest that the object category information might be more accessible in specific time spans from around 50 ms to 300 ms in the EEG signals. However, it has not been shown how (if at all), other features of the EEG signals (i.e. all except Mean), reflect object category information across the time course of trials. To answer this question, we split the trials into six 100ms sub-windows of time and evaluated the category information within each sub window from each individual feature (Fig. 5). Please note that, by definition, we excluded the ERP components in this analysis. The results showed that the highest information was observed in the 100-200 ms (for almost all features except fractal dimensions, approximate entropy and SEF) followed by the 200-300 ms sub windows. Interestingly, rather the commonly used signal Mean feature, the maximum average decoding accuracies were obtained from the within-trial correlation and the Hurst exponent both of which measure aspects of signal autocorrelation in the EEG. While these results are consistent with previous observations showing category-related information in the 50 ms to 300 ms time windows (Karimi-Rouzbahani et al., 2019; Karimi-Rouzbahani, 2018), they also suggest that features of EEG signals, which detect temporally correlated patterns (i.e. within-trial correlation and Hurst exponent), can provide additional category-related information for decoding.

**Fig. 4.**
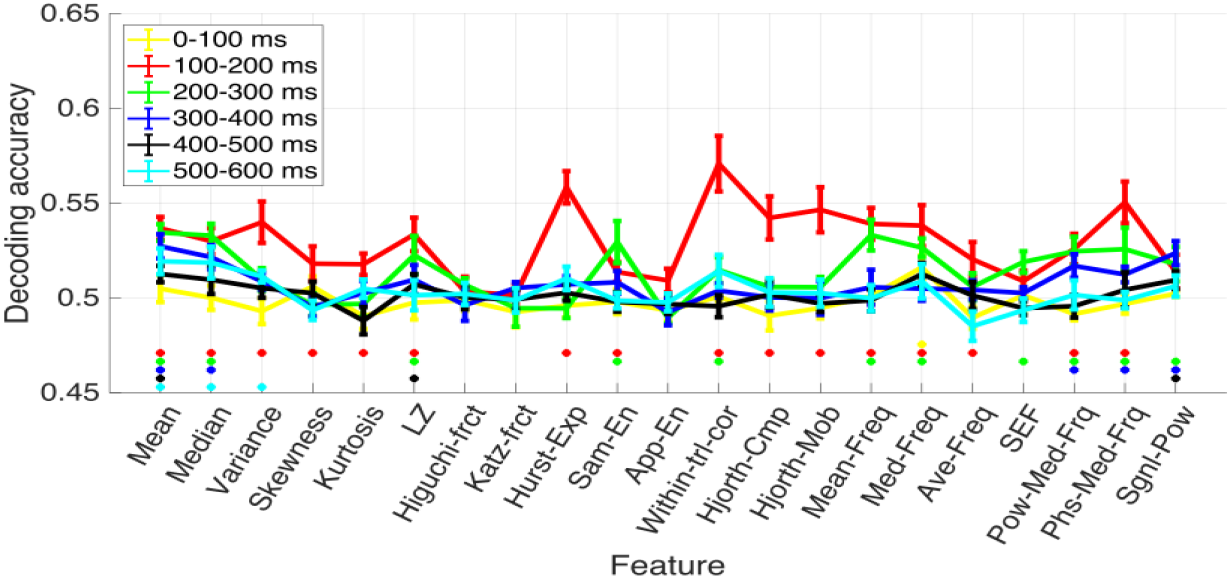
Decoding accuracy obtained for all features (except the time-dependent ERP features) in six 100ms sub windows of trials. Decoding results reflect the average classification accuracy along with the standard error (SE) across participants (error bars). Colored dots indicate features with significantly (p<0.05; random permutation testing) above-chance accuracy.

## 4 Discussion

While the artificial intelligence-based object recognition algorithms have started a remarkable progress after the introduction of the brain-plausible convolutional neural networks (Krizhevsky et al., 2012), we are still short of enough understanding about how the brain processes categories of objects for recognition. This understanding is critical for improving the AI-based algorithms. We believe that the first step in understanding the human brain is to find the optimal way in which we can read out/decode the information processing of the brain. In this work, we approached this question by evaluating the information content of a large set of features extracted from human brain activity (EEG) during object recognition. We observed that, the well-established ERP components of the EEG were the most informative features in the feature set and their information was more pronounced in the Theta band rather than any other bands. Upon limiting the analysis time window to 100 ms windows, we observed that the information was mainly concentrated in the 100-200 ms time window (at around the same time of the ERPs) and it was more pronounced for the features which detected repetitive patterns across time (i.e. within-trial correlation and Hurst exponent). Finally, we showed that the information about human faces were more accessible in the brain activity.

These results have several implications for future developments of brain-inspired object recognition algorithms. First, the information about object category processing should be looked for in the span from 50 ms to 1000 ms post stimulus onset, when trying to study the human object recognition brain. This could confine many previous work which looked at the whole trial time window for additional category-related information (Taghizadeh-Sarabi et al., 2014; Torabi et al., 2017) and can inform future studies. Second, brain decoding for object category information should look at the Theta band as it is the main substrate for the coding of feed-forward visual processing of visual information in the brain (Bastos et al., 2015). Third, temporal variability of brain activity should not be overlooked, as this is generally the case in multivariate decoding (Karimi-Rouzbahani et al., 2019), as the temporal codes can contain additional information. Forth, as the brain seem to process the face and object information using a common set of neural infrastructures (Dobs et al., 2019), but constructs face representations which outperform object representations (Fig. 2B), the current deep networks of object recognition, may benefit to follow the brain and move towards a unified platform for face and object recognition. This is not the case as we generally use separate deep neural networks for the categorization of objects (e.g. AlexNet) and faces (e.g. VGG face).

This work also provides new insights for the research in the area of Brain-Computer Interface (BCI), by putting the spotlight on the most informative features that can improve the classification accuracy. Please note that we did not try to maximize the decoding accuracy in this study, but rather, tried to see how the brain processes information about object categories. That is why we used unsupervised feature extraction methods to see how the brain activity reflects information, rather than using supervised methods for feature transformation and combination which are used in the BCI domain (Pouryazdian & Erfanian, 2009). Of course, using supervised machine learning methods can potentially provide advantageous decoding accuracies on this same dataset.

Together, this paper presents preliminary results about where we should look for object category information when decoding brain activity. There can be many interesting future directions for this work including combining the informative features (Rouzbahani & Daliri, 2011), fusing the neural representations of objects (EEG) with those of state-of-the-art convolutional neural networks to see their similarities/differences using representational similarity analysis (Hamid Karimi-Rouzbahani, 2018), etc.

